# Mechanical stretch sustains myofibroblast phenotype and function in microtissues through latent TGF-β1 activation

**DOI:** 10.1101/2020.01.25.917179

**Authors:** Matthew Walker, Michel Godin, Andrew E. Pelling

## Abstract

Fibrosis is a leading cause of death in developed countries that is characterized by a progressive deterioration of tissue mechanical behavior. Developing methods to study tissue mechanics and myofibroblast activation may lead to new targets for therapeutic treatments that are urgently needed. Microtissue arrays are a promising approach to conduct relatively high throughput research into fibrosis as they recapitulate key biomechanical aspects of the disease through a relevant 3D extracellular environment. In early work, our group developed a device called the MVAS-force to stretch microtissues while enabling simultaneous assessment of their dynamic mechanical behavior. Here we investigated TGF-β1 induced fibroblast to myofibroblast differentiation in microtissue cultures using our MVAS-force device through assessing α-SMA expression, contractility and stiffness. By doing so, we linked cell-level phenotypic changes to functional changes that characterize the clinical manifestation of fibrotic disease. As expected, TGF-β1 treatment promoted a myofibroblastic phenotype and microtissues became stiffer and possessed increased contractility. Furthermore, these changes were partially reversible upon TGF-β1 withdrawal. In contrast, however, long-term cyclic stretching maintained myofibroblast activation. Furthermore stretching had no effect compared static cultures when TGF-β1 receptors were inhibited and stretching promoted myofibroblast differentiation when given latent TGF-β1. Together these results suggest that external mechanical stretch may activate latent TGF-β1 and might be a powerful stimulus for continued myofibroblast activation to progress fibrosis. Further exploration of this pathway with our approach may yield new insights into myofibroblast activation and more effective therapeutic treatments for fibrosis.

**Insight box:** Using a novel high-throughput approach, we quantified the effects of dynamic mechanical stretching on the phenotype and function of cells in 3D microtissue cultures during myofibroblast activation with TGF-β1 treatment and subsequent withdrawal. Our findings show that mechanical stretch may activate endogenously produced latent TGF-β1 to maintain the presence and activity of myofibroblasts after tissue injury. Importantly, through this feed forward mechanism, mechanical stretch might be a powerful stimulus that directs tissues away from recovery and towards the development of fibrosis.

## Introduction

Myofibroblast activation is a normal healing response following tissue injury found throughout the body, and is essential for rapid wound contraction and *de novo* matrix deposition^1–5^. However, when unchecked, continued myofibroblast activation may lead to chronic fibrosis, and potentially, a life-threatening loss of tissue functionality^1, 6, 7^. As one of the leading causes of death in developed countries, fibrosis has become an area of high interest for lung, heart, vasculature, liver, renal, and eye research^4, 8–11^. Yet the contributing factors that determine whether myofibroblast activation persists and fibrosis progresses into a chronic illness, or whether restoration occurs and tissues regain their initial functionality, remain unclear^4, 11^. Identifying these factors is not only necessary to understand how the activation of myofibroblasts is controlled, but importantly, may also provide valuable insights for future therapeutic approaches to prevent the development and arrest the progression of fibrosis.

Transforming growth factor (TGF)-β1, a biochemical inflammatory mediator, has been identified as a key determinant in the development and progression of tissue fibrosis, and as such, has become a central focus of research^12, 13^. In that regard, TGF-β1 treatment is routinely used to induce fibroblast to myofibroblast differentiation *in vitro* in lieu of any other biochemical stimuli^14, 15^, and overexpression of TGF-β1 in animal models consistently exhibit marked fibrotic changes^16, 17^. It is well recognized that TGF-β1 is a potent agonist of the SMAD2/3 pathway^14^, which in fibroblasts leads to α-smooth muscle actin (α-SMA) expression, a commonly used biomarker of myofibroblast differentiation^2, 4, 5^. In addition to canonical SMAD2/3 signaling, TGF-β1 may also activate mitogen-activated protein kinase (MAPK) pathways to further regulate differentiation, proliferation, cell survival and apoptosis^18^.

In an autocrine feedforward loop to drive further myofibroblast activation, inactivated TGF-β1 is secreted by myofibroblasts as a large latent complex (LLC) consisting of the latent TGF-β1 binding protein (LTBP), which associates with the extracellular matrix (ECM), and the latency-associated peptide (LAP), which non-covalently sequesters a TGF-β1 polypeptide and binds to integrins on the cell’s membrane^19–21^. Mechanical stretch, such as encountered in lung tissue from breathing or internally produced by myosin contraction, has been previously proposed as a mechanism to activate this latent source by directly inducing a conformational change in the LAP to release active TGF-β1 into the ECM^19, 22–24^. In doing so, this mechanosensitive pathway may maintain myofibroblast activation to advance the tissue towards of chronic fibrosis.

Although fibrosis and therapeutic treatments have often been investigated *in vitro* to reduce complexity and increased control, the effectiveness of this approach is often hindered by the lack of a biochemically and mechanically relevant environment when using standard 2D cell culture techniques^25, 26^. Instead microtissue arrays, in which cells self-assemble within a collagen ECM around vertical cantilevers into sub-millimeter 3D organized structures, offer an appealing high-throughput alternative^27, 28^. In addition to a relevant ECM, the microtissue method also enables direct assessment of tissue-level functional changes through tracking the visible deflection of the cantilevers for contractility measurements. Because these abilities, this method has been shown to be especially appropriate for testing the efficacy of fibrosis treatments, as the clinical manifestation of fibrosis is stiff, chronically contracted tissues with a remodeled ECM^28^.

In previous work, we developed a microtissue vacuum actuated stretcher (MVAS-force) that was capable of stretching an array of microtissues and simultaneous measurements of dynamic stiffness and contractility^29^. Here we used our MVAS-force device to link changes in α-SMA expression during TGF-β1 treatment to mechanical changes that are characteristic of tissue fibrosis. We then assessed whether microtissue mechanics and the quiescent fibroblastic phenotype is restored following TGF-β1 withdrawal. Finally, we tested the hypothesis that stretch may maintain myofibroblast differentiation during TGF-β1 withdrawal through endogenously produced latent TGF-β1 activation. These findings contribute to the field’s understanding of the progression of fibrosis through TGF-β1 induced fibroblast to myofibroblast differentiation. We also established that MVAS-force devices are capable of investigating fibrosis development, and as such, may be a useful tool for pharmacological discovery. Lastly our findings further implicate that mechanical forces are stimuli that direct tissues away from normal healing pathways and towards fibrotic development, which may have important implications in therapeutic treatment development.

## Methods

### Cell culture

Prior to microtissue fabrication, NIH3T3 (ATCC) fibroblast cells were maintained on 100mm tissue culture dishes (Fisher) at 37℃ with 5% CO_2_ until 80-90% confluent in feeder media composed of Dulbecco’s Modified Eagle’s Medium (DMEM) supplemented with 10% fetal bovine serum (FBS), 100mg/ml streptomycin and 100U/ml penicillin antibiotics (all from Hyclone Laboratories Inc.).

### Microtissue fabrication

MVAS-force devices consist of 6 rows of 10 wells, each containing a microtissue formed around a pair of vertical cantilevers. To stretch the microtissues, vacuum chambers border each row, and are connected to independent external regulators (SMC ITV0010) controlled through Labview. When a vacuum is applied, the cantilevers closest to the vacuum chamber are actuated, while tissue tension can be measured through the visible deflection in the opposing ‘force-sensing’ cantilevers.

MVAS-force devices were fabricated from mold replication steps described previously^29, 30^. Briefly, SU-8 masters for the three layers of the device were created with standard photolithographic techniques. The top layer contains the cell culture wells and vacuum chambers, the middle layer is a thin membrane with the cantilevers, and the bottom layer includes vacuum and empty bottom chambers. To fabricate devices, masters were cast with polydimethylsiloxane (PDMS) containing a 1:10 cross linker ratio and plasma bonded together.

Microtissues were cultured in MVAS-force devices as described previously^27, 29, 30^. Prior to seeding, devices were sterilized with 70% ethanol and treated with 0.2% Pluronic F-127 (P6866, Invitrogen) for two minutes to reduce cell adhesion. ∼650 cells were centrifuged into each well in a solution containing 1.5mg/ml rat tail collagen type I (354249, Corning), 1x DMEM (SH30003.02, Hyclone), 44 mM NaHCO_3_, 15 mM d-ribose (R9629, Sigma Aldrich), 1% FBS and 1 M NaOH to achieve a final pH of 7.0-7.4. Excess collagen was removed and devices were incubated at 37°C for 15min to initiate collagen polymerization. Lastly, an additional ∼130 cells were then centrifuged on top of each tissue. Feeder media was added and changed every 24 hours. Microtissues were allowed to compact and secure themselves to the cantilevers under static conditions with feeder media for 2 days prior to experimentation.

### Myofibroblast differentiation

To assess myofibroblast differentiation, the cell culture media was switched to a differentiation formulation containing DMEM, 5% FBS, 100mg/ml streptomycin, 100U/ml penicillin antibiotics, and 5ng/ml TGF-β1 (Peprotech, 100-21). Functional assessment of differentiation was assessed every 24 hours over a 3-day period. Because there was no change to microtissues beyond day 2, differentiation for subsequent experiments was kept to 2 days. To block the differentiation and to assess the contribution of Rho-signaling, the differentiation media was supplemented with 1µM Y27632 (Y27) (Y0503, Sigma Aldrich), a rock inhibitor.

We next investigated whether myofibroblast differentiation was reversible, by switching microtissues back to feeder media for 2 days following 2 days of differentiation. Once it was demonstrated that microtissue differentiation was in part reversible, we tested the hypothesis that long-term oscillatory mechanical stretch may maintain myofibroblast differentiation by cyclically loading microtissues at 0.5Hz 5% strain during dedifferentiation. Further in regards to this hypothesis, to assess whether stretch activates endogenously produced TGF-β1, TGF-β1 receptors were blocked with 10µM GW788388 (GW) (16255-1, Cayman Chemicals) during dedifferentiation, and lastly, differentiation was assessed with a latent source of 5ng/ml TGF-β1 (299-LT-005, R&D Systems) under both static and stretching conditions.

### Functional assessment of differentiation

To assess how microtissues are functionally altered during myofibroblast differentiation, microtissue mechanics were assessed *in situ* using the MVAS-force at 37°C and under 5% CO_2_^29^. Briefly, forces were measured through the visible deflection of the force-sensing cantilever and its known spring constant. To calculate the deflection, images were captured at focal planes containing both the top and bottom of the cantilevers. The bottom position of the cantilevers was measured using a centroid algorithm while the top was tracked using pattern matching with adaptive template learning in Labview. Static deflections, Τ_o_, were calculated by subtracting the top and bottom positions and gave an assessment of the resting contractility.

To assess stiffness changes, the dynamic mechanical behaviors were measured at 0.5Hz with 3% strain. Images at both the tops and bottoms of the cantilevers were captured at 15fps for a minute. Tissue tension was calculated from the difference in the positions of the force-sensing cantilever in those images after accounting for the phase shift caused by the camera delay between capturing the two focal planes. Microtissue strain, ε, was defined as the percent change in length measured from the innermost edges of the tops of the cantilevers (equation 1). The storage stiffness, a measurement of elasticity, was then calculated as the ratio of the magnitudes of the Fourier transforms of tension and strain multiplied by the cosine of the phase lag between force and strain (equation 2).

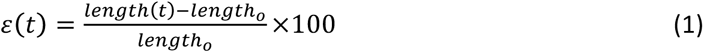

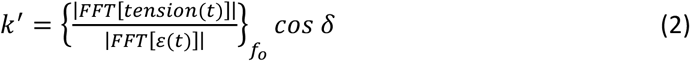

### Phenotypic assessment of differentiation

α-SMA was used as biomarker^2, 4, 5^ to assess myofibroblast differentiation, and imaged with standard immunofluorescence techniques. Briefly microtissues were fixed and permeabilized *in situ* with ice-cold methanol for 10min. To prevent non-specific binding, microtissues were blocked for 30min with 5% FBS. α-SMA was labeled with 1:100 primary antibody produced in rabbit (Abcam, ab5694) at room temperature for two hours and then 1:200 Goat anti-rabbit IgG secondary antibody conjugated to Alexa Fluor 488 (Invitrogen, A11034) at room temperature for an addition two hours. Cell nuclei were stained with DAPI (Fisher, D1306).

Image stacks of microtissues were acquired on a TiE A1-R laser scanning confocal microscope (LSCM) (Nikon) with appropriate laser lines and filter blocks. To produce heat maps, images were flattened by integration, spatially aligned and averaged together. α-SMA expression was normalized to the nuclei fluorescence to give a per cell measurement and to account for differences in proliferation.

### Data analysis and statistics

All numerical data are presented as mean ± standard error. Statistical tests, as described in the results, were performed using Originlab 8.5 (Northampton, MA) with p<0.05 considered statistically significant.

## Results

### Myofibroblast differentiation in microtissue cultures

Myofibroblast differentiation has historically been characterized through biochemical assays of biomarker expression (ie. α-SMA)^2, 4, 5^ in cells grown in 2D culture. Comparably there have been much fewer *in vitro* investigations that have linked the appearance of a myofibroblastic phenotype to functional changes in mechanical behavior (ie. increased stiffness and contractility) at the tissue-level which are characteristic of fibrosis^28, 31^. For that reason, we assessed both changes to static contractility (fig.1a) and dynamic stiffness (fig. 1b) of microtissues during myofibroblast differentiation using our MVAS-force device.

**Figure 1:**
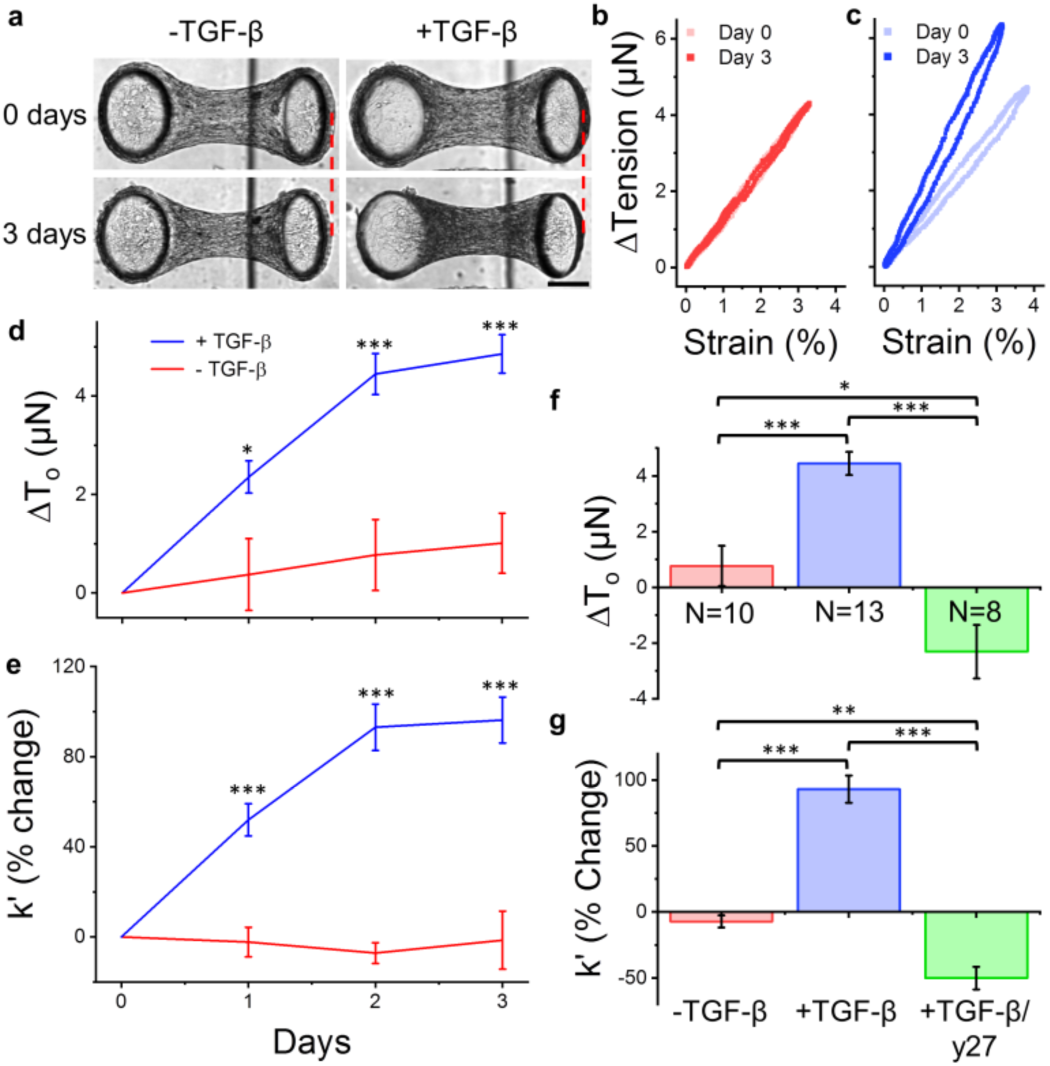
TGF-β1 treatment contracts and stiffens microtissue cultures. At experimental day 0, once microtissues had compacted and were securely anchored to the cantilevers (2 days after seeding), they were either switched to a media containing TGF-β1 to promote myofibroblast differentiation or kept in normal feeder for an additional 3 days. TGF-β1 treatment increased the resting tension, Τ_o_, compared to feeder media indicated by a greater deflection in the force-sensing cantilever (a). Additionally the dynamic stiffness of microtissues was measured *in situ* with the MVAS-force at 0.5Hz. Representative clockwise oriented tension-strain loops with normal feeder and differentiation media are in (b) and (c), respectively. While there was no difference in stiffness between experimental days 0 and 3 with feeder media, the stiffness markedly increased with TGF-β1 treatment. The average changes to the resting tension and stiffness over the 3-day period are in (d) and (e), respectively. The TGF-β1 induced increased tension and stiffness largely occurred over the first 2 days, leveling off by day 3. The effects of TGF-β1 treatment were completely blocked with concurrent treatment with y27, a ROCK inhibitor, indicating the role of Rho-induced stress fiber formation and myosin contraction during myofibroblast differentiation (f and g). The scale bar in (a) represents 100µm. *P<0.05, **P<0.01, ***P<0.001.

As expected, TGF-β1 treatment increased the resting tension and stiffness over a 3-day period compared to the control group (fig. 1d and e, respectively) (t-tests). Functional changes largely occurred over the first 2 days of treatment, with no significant difference to either contractility or stiffness between days 2 and 3 (repeated measures t-tests p>0.05). For this reason, 2 days was used as the time point for all subsequent experiments. After 2 days of TGF-β1 treatment, there was a 4.4 ± 0.4µN increase in contractility and a 93 ± 10 % increase in stiffness.

Inhibiting Rho-signaling with the ROCK inhibitor y27 blocked TGF-β1 induced functional changes (fig. 1f and g). In fact, concurrent treatment of TGF-β1 with y27 reduced microtissue contractility and stiffness below control microtissues maintained in feeder media (P<0.05 and <0.01, 1-way ANOVA). These results are not overly surprising as Rho is known to direct the assembly and stabilization of the actin cytoskeleton^32^ and control contractility through deactivation of myosin phosphatase and phosphorylation of myosin light chain^33^. Several studies have also shown the importance of Rho/ROCK signaling in controlling collagen synthesis^34–36^, which would also contribute to increased microtissue stiffness.

To link functional changes in microtissues to a myofibroblast phenotype change, α-SMA was immunofluorescently imaged. Average heat maps of the distribution of cells and α-SMA expression indicated that TGF-β1 treatment lead to myofibroblast differentiation predominately in cells directly associating with the relatively stiff PDMS cantilevers and around the perimeter of the tissue (fig. 2a). Moreover, the distribution of cells in the tissues became less uniform as TGF-β1 treatment increased the concentration of cells towards the center of the tissue. In contrast, however, there was no change to the total microtissue width on bright field images (data not shown, P>0.05, T-test). The spatial distributions were not greatly affected with concurrent treatment of TGF-β1 and y27 compared to TGF-β1 alone.

**Figure 2:**
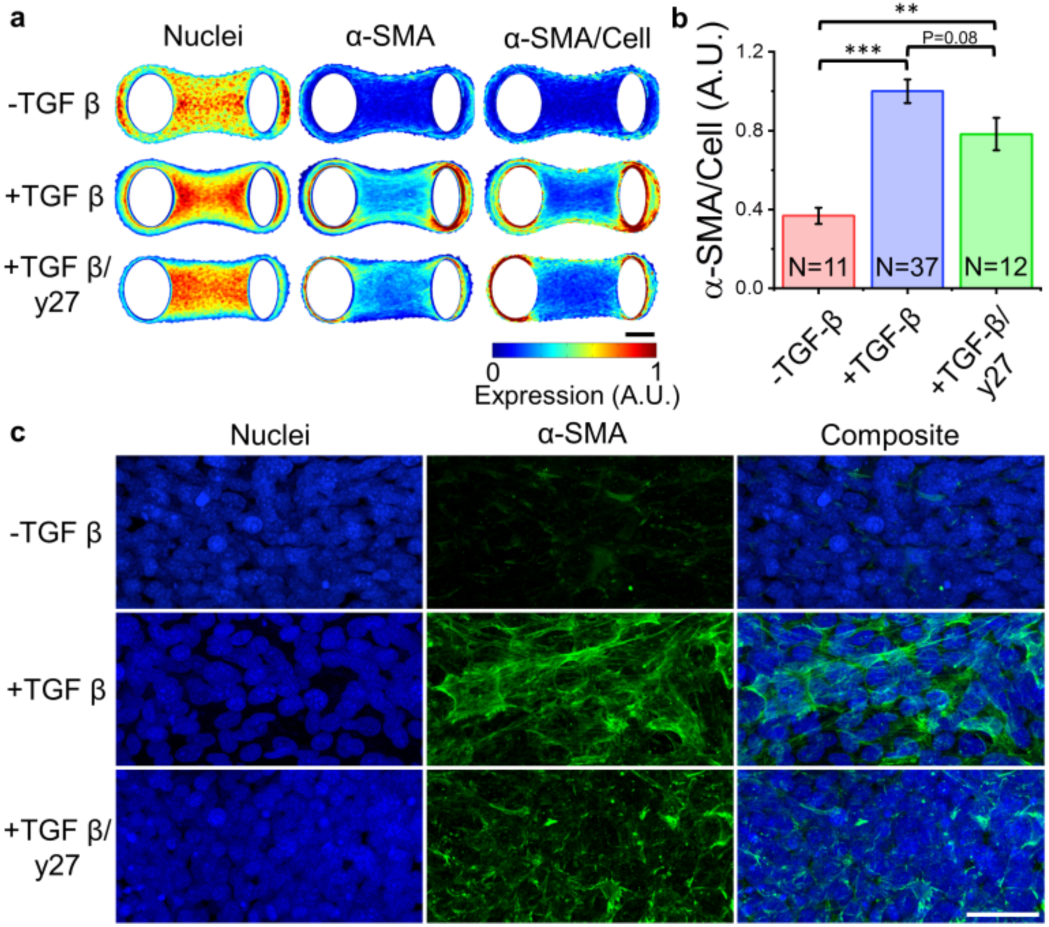
TGF-β1 induces myofibroblast differentiation of microtissues. To quantify TGF-β1 induced myofibroblast differentiation, microtissues were stained for α-SMA. Average heat maps of cell nuclei, α-SMA expression and normalized expression are in (a). Total α-SMA expression normalized to the number of cells is in (b). TGF-β1 treatment significantly increased α-SMA expression compared to the control. Concurrent treatment with y27 still had greater α-SMA expression than control but slightly, albeit not significantly, less than TGF-β1 alone. Images of representative centrally located magnified regions are in (c). TGF-β1 treatment promoted the expression and polymerization of dense α-SMA fibers. In microtissues that concomitantly received y27, α-SMA was not as densely polymerized into organized fibers, indicating the role of Rho in the recruitment of α-SMA into stress fibers. The scale bars in (a) and (c) represent 100 and 50µm, respectively. **P<0.01, ***P<0.001.

Overall, the total normalized α-SMA expression increased by 120% with TGF-β1 treatment compared to the control (fig. 2b) (1-way ANOVA, p<0.001). α-SMA expression was also increased with concurrent treatment of TGF-β1 and y27 (P<0.01). However, compared to TGF-β1 alone, y27 treatment seemed to decreased α-SMA expression, albeit the difference was not statistically significant (P=0.08). In representative images of centrally located regions, α-SMA is organized into polymerized fibers with TGF-β1 treatment (fig. 2c). Whereas in microtissues that concomitantly received y27, α-SMA was less densely polymerized into organized fibers. These findings demonstrate the involvement of Rho/ROCK pathways in the recruitment of α-SMA into stress fibers.

### Mechanical stretch sustains myofibroblast differentiation

The initial appearance of myofibroblasts is a normal physiological response following injury, contributing to wound contraction and collagen matrix deposition^1–5^. In tissues that are fully repaired, the myofibroblast phenotype disappears from the wound space and tissues regain their initial functionality. For reasons unknown, however, the presence of myofibroblasts may linger beyond normal wound healing, leading to a loss of tissue functionality and chronic fibrosis^1, 6, 7^. Accordingly we assessed whether microtissues may recover their initial fibroblastic phenotype and functionality if the TGF-β1 treatment was withdrawn following 2-days of differentiation.

Switching microtissues back to feeder media for 2 days decreased contractility by −2.1±0.3µN and stiffness by −18 ± 3% (1-way ANOVA, P<0.01 and P<0.001) (fig. 3a and b). Removing TGF-β1 also reduced α-SMA expression by 40 ± 4% (1-way ANOVA, P<0.001) (fig. 3c,d,e). Importantly these findings show that the presence and functionality of myofibroblasts can in part be reversed in microtissues that were differentiated as in a normal wound healing process.

**Figure 3:**
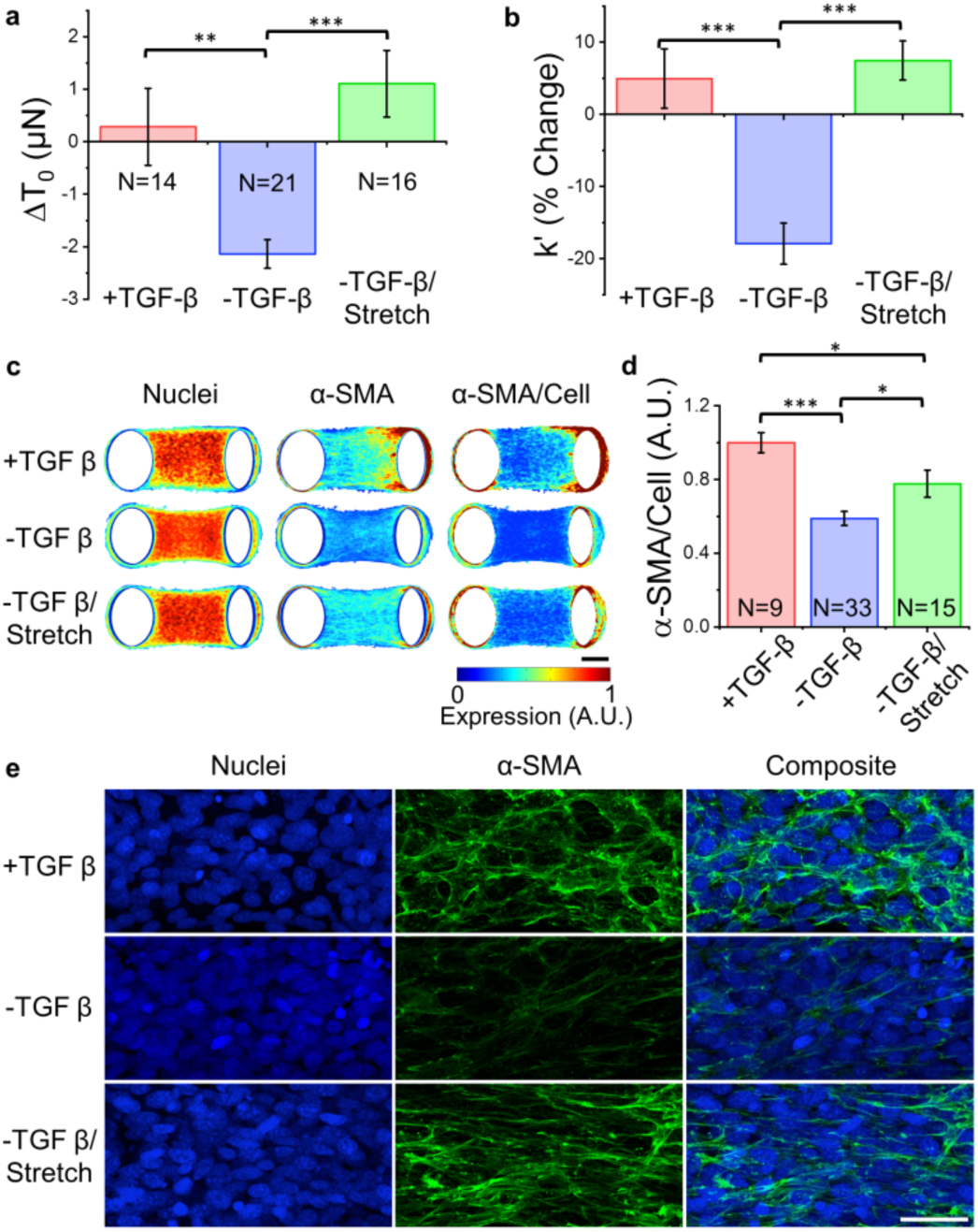
Myofibroblast differentiation is, in part, reversible but sustained through mechanical stretch. To assess whether the activation of the myofibroblast phenotype could be reversed, microtissues were allowed to compact for 2 days in feeder media, then differentiated with TGF-β1 media for 2 days and finally switched back to feeder media. Compared to microtissues that were sustained in differentiation media, TGF-β1 withdrawal decreased resting tension and stiffness (a and b, respectively). In contrast, however, the tension and stiffness did not decrease in microtissues that were dynamically stretched at 5% 0.5Hz in addition to TGF-β1 withdrawal. These changes were mirrored in heat maps, normalized total, and representative images of α-SMA expression (c-e, respectively). TGF-β1 withdrawal decreased α-SMA expression and fiber formation. In contrast dynamic stretching, in part, prevented the loss of myofibroblast phenotype, but was still significantly reduced compared to continued TGF-β1 treated microtissues. These findings indicate that mechanical stretch may sustain myofibroblast differentiation. The scale bars in (c) and (e) represent 100 and 50µm, respectively. *P<0.05, **P<0.01, ***P<0.001.

Mechanosensing pathways have been previously identified as key stimuli in myofibroblast differentiation^3, 5, 37, 38^, however, the role of dynamic stretch remains unclear. In that regard, stretching microtissues during TGF-β1 treatment produced no functional or phenotypic differences (SI 1). In contrast, however, stretching microtissues upon TGF-β1 withdrawal prevented the decrease in contractility and stiffness that had been observed in static cultures that were also switched back to feeder media. In fact, stretching with TGF-β1 withdrawal sustained microtissue mechanics to the same extent as continued TGF-β1 treatment (P>0.05). This maintenance of microtissue mechanics was also mirrored by an increased α-SMA expression compared to static conditions (1-way ANOVA P<0.05). Admittedly, however, the expression was still lower than in microtissues kept in TGF-β1 (P<0.05). Nevertheless, these results suggest that cyclic stretching may in part be responsible for the maintenance of myofibroblastic phenotype, leading to chronic fibrosis.

### Mechanical stretch activates latent TGF-β1 in microtissues

It has been previously hypothesized that dynamic stretching may contribute to myofibroblast differentiation through activating latent TGF-β1 sources sequestered by matrix proteins. This latent form can be produced and secreted into the extracellular environment by myofibroblasts in a feedforward loop, to drive further differentiation^19–21^. Therefore to assess whether mechanical stretching sustained myofibroblast differentiation through activating endogenous TGF-β1, we attempted to dedifferentiate microtissues while TGF-β1 receptors were blocked with GW to inhibit autocrine signaling.

As expected, blocking receptors did not affect the observed decrease to the stiffness or α-SMA expression compared to TGF-β1 withdrawal only (1-way ANOVA, p>0.05) (fig 4a and c). Furthermore, when receptors were blocked, microtissue stiffness and α-SMA expression, distribution and organization were no different under static and stretching conditions (1-way ANOVA, p>0.05) (fig. 4 a, b, c and d). These results indicate that stretch maintains myofibroblast functionality and phenotype through an autocrine TGF-β1 signaling.

**Figure 4:**
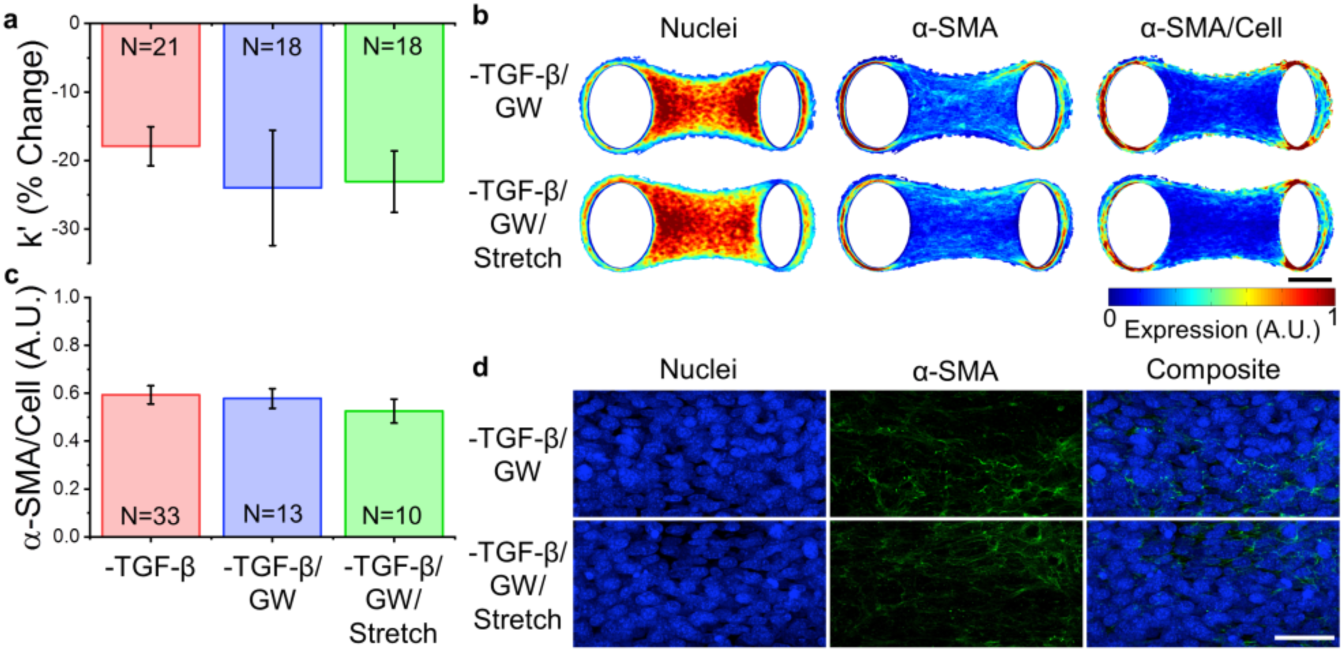
TGF-β1 receptor inhibition prevents sustained myofibroblast differentiation through mechanical stretch. To assess whether dynamic stretching sustained myofibroblast differentiation through activation of endogenous latent TGF-β1 in an autocrine signaling pathway, TGF-β1 receptors were blocked with GW during exogenous TGF-β1 withdrawal. Upon blocking TGF-β1 receptors, dynamic stretching did not effect the reduction of microtissue stiffness (a) or α-SMA expression (b-d) that accompanied exogenous TGF-β1 withdrawal. These findings suggest that the sustained differentiation with mechanical stretching is through endogenous TGF-β1 activation. The scale bars in (b) and (d) represent 100 and 50µm, respectively.

In the latent form, TGF-β1 is non-covalently caged by the mechanosensitive latency-associated peptide (LAP), which effective decreases its bioavailability. It has been hypothesized that stretch may cause conformational changes to the LAP to free TGF-β1 to promote continued myofibroblast differentiation. Therefore to assess whether dynamic stretch was capable of causing this conformational change, microtissues were differentiated with 5ng/ml Latent TGF-β1 for 2 days.

Latent TGF-β1 alone produced no change in the stiffness (repeated measures t-test, p>0.05) and α-SMA expression remained low throughout microtissues (fig. 5). In contrast, dynamic stretch, in addition to latent TGF-β1, increased microtissue stiffness and α-SMA expression (t-tests, P<0.001). Admittedly, the stiffness and expression levels still remained lower than if the same concentration of activated TGF-β1 was given, likely because stretch did not activate 100% of the latent TGF-β1 given to the microtissues. Nevertheless, these results indicate that stretch can mechanically activate latent TGF-β1.

**Figure 5:**
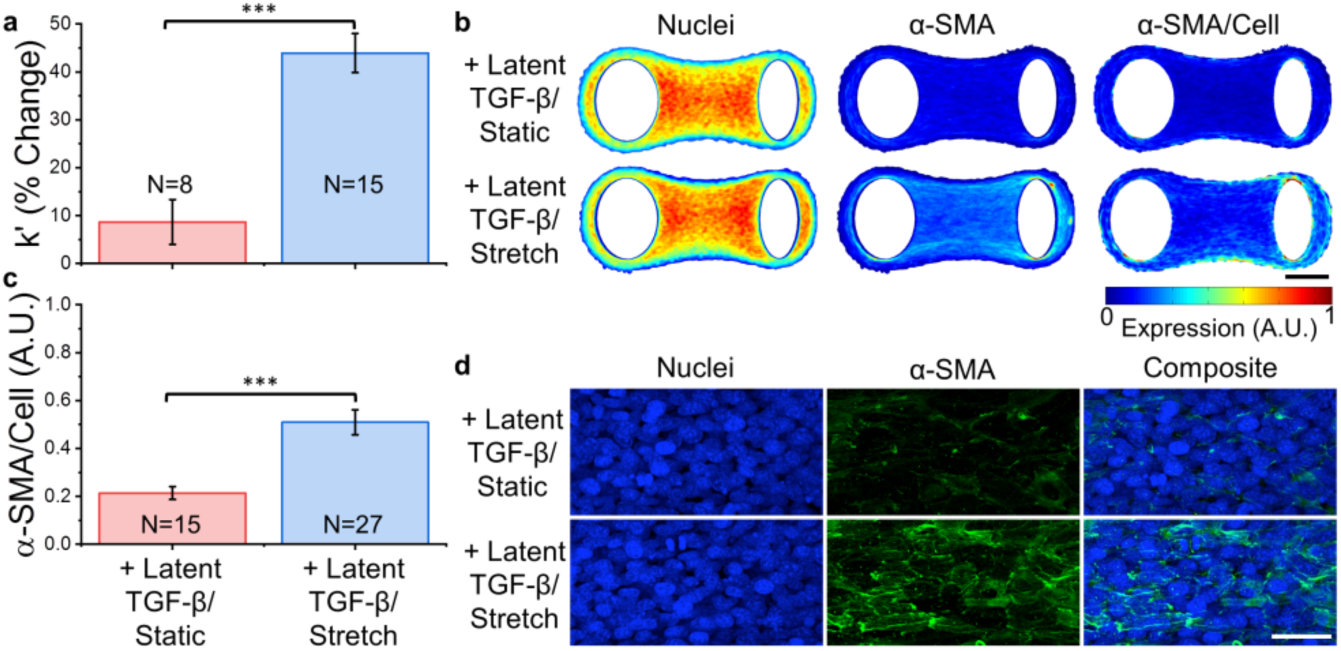
Latent TGF-β1 is activated by mechanical stretch to promote myofibroblast differentiation. To demonstrate that a latent source of TGF-β1 can be activated by mechanical stretch, myofibroblast differentiation was assessed under static and stretching conditions with exogenous latent TGF-β1. Dynamic stretch significantly increased microtissue stiffness (a) and α-SMA expression (c-d) compared to static conditions. These findings indicate that mechanical stretching can activate latent TGF-β1, further supporting its role in sustaining myofibroblast differentiation during TGF-β1 withdrawal. The scale bars in (b) and (d) represent 100 and 50µm, respectively. ***P<0.001.

## Discussion

Continued research and development of techniques are urgently needed to further our understanding of myofibroblast activation and to discover new drug targets to treat fibrosis. Not only do microtissues recapitulate key biochemical and mechanical properties of both healthy and fibrotic tissues, they allow cell-level phenotypic and tissue-level functional screening in a relatively high-throughput manner. For these reasons, microtissues are a compelling approach to conduct fibrosis research^28^. Here we aimed to investigate myofibroblast activation and deactivation under cyclic stretching in a microtissue model using our MVAS-force device. Importantly this approach allowed us to link changes to protein expression to mechanical behaviors that directly reflect the clinical manifestation of fibrotic disease.

As expected, TGF-β1 treatment led to fibroblast to myofibroblast differentiation, indicated by α-SMA expression, and increased microtissue contractility and stiffness. These changes were consistent with a previous publication that assessed microtissues populated with human lung fibroblasts^28^. Our findings are further in accordance with multiple studies have shown that increased tissue stiffness is a characteristic feature of fibrosis, albeit, the changes we observed over a relatively short experimental time were less than those previously reported in fibrotic lungs (increases from ∼2kPa to ∼17kPa)^39–41^ and liver (∼1kpa to 3-22kPa)^42^. In organs with established fibrosis, stiffness changes are thought to arise from extracellular protein deposition, particularly fibrillar collagen, and matrix crosslinking^42, 43^. Increased deposition of collagen fibers upon myofibroblast activation has already been reported in microtissues^28^, and thus, likely contributed to our results. Altered cell (as opposed to matrix) stiffness may also have made a significant contribution to the change in tissue mechanics. In that regard, it has been previous argued that differences between decellularized matrices cannot fully account for differences in stiffness between normal and fibrotic tissue samples^39^. Although the contribution from altered cell properties to fibrotic tissue stiffness has yet to be established, it may be considerable^7^.

A stiffer ECM is well accepted to encourage myofibroblast activation through mechanosensing at focal adhesions and Rho/ROCK signaling^37, 38^. Therefore, it was not surprising that we found that concurrent ROCK inhibition during TGF-β1 treatment prevented fibrotic functional changes and helped preserve the fibroblastic phenotype. Accordingly the Rho/ROCK pathway has become an attractive target for anti-fibrotic pharmaceutical treatments^44–46^, and from our demonstration here, the MVAS-force may offer a method for testing and developing such treatments in tissue-like structures prior to clinical trials.

In addition to mechanosensing through focal adhesions, α-SMA itself is beginning to be considered mechanosensitive, as it only localizes to stress fibers when cells are subjected to considerable mechanical loads^3^. For example, α-SMA expression in stress fibers requires a substrate stiffness around 20kPa and reducing cellular tension by inhibiting myosin contraction leads to disassembly of α-SMA stress fibers^47, 48^. This could provide a direct explanation as to the tight coupling between tissue mechanics and α-SMA expression that we generally observed outside of ROCK inhibition.

Although it is generally agreed that increased substrate stiffness favors fibroblast differentiation into myofibroblasts^49, 50^, the role of cyclic mechanical stretch, as cells would encounter in airways and vasculature, is less clear. There have been conflicting reports as to whether cyclic stretch promotes myofibroblast differentiation^51–53^ or serves as a protective role promoting a quiescent fibroblastic phenotype^54, 55^. In contrast to those reports, we did not find that stretch had any effect on myofibroblast activation in microtissues. Differences in ECM stiffness may partially explain these conflicting reports, since stretching of lung fibroblasts cultured on stiff (30kPa) gels has been shown to augment myofibroblastic phenotype, while stretching on soft (2kPa) gels had no affect^53^. Differences in the strain field may also contribute as enhanced anisotropy when under biaxial loading has been shown to augment myofibroblast activation from valve interstitial cells^52^.

Following stimulation with TGF-β1, subsequent withdrawal partially reversed the myofibroblastic expression phenotype and mechanical behavior. Classically in the paradigm for tissue repair, terminally differentiated myofibroblasts are removed from healing tissue through apoptosis^56^. However, opposed to earlier investigations which suggested that TGF-β1 induces a relatively stable alteration in cell phenotype^14^, more recent work has found that myofibroblasts have the capacity for de-differentiation back into a quiescent fibroblast phenotype^31, 57–59^.

Importantly in contrast to microtissues kept under static conditions, we found that cyclic stretching maintained the myofibroblastic phenotype and tissue mechanics during TGF-β1 withdrawal. This can likely be accounted by autocrine signaling involving mechanical activation of latent TGF-β1 since blocking TGF-β1 receptors while stretching led to identical behaviors as in static conditions. We further demonstrated that stretching microtissues could activate latent TGF-β1 to promote differentiation towards a myofibroblast phenotype and to increase tissue stiffness.

These results add to an already large precedent that TGF-β1 can be mechanically activated from extracellular stores. For example, contractile forces from lung fibroblasts have been shown to directly activate latent TGF-β1 from ECM independent of proteolytic activity^19^. Furthermore, tensile loading of fibrotic lung strips respond by releasing active TGF-β1, and subsequently increasing Smad2/3 signaling^22^. Lastly, using single molecule force spectroscopy to pull directly on LAP has been shown to induce a conformation change that frees active TGF-β1 in a pure mechanical process^23, 24^. Admittedly we found no difference in deactivation with and without a TGF-β1 receptor block under static conditions, and exogenous latent TGF-β1 treatment had no effect on static cultures. Together these results indicate that endogenous latent TGF-β1 did not contribute to responses under static culturing. Although contractile forces produced by myofibroblasts during wound contraction are thought to be strong enough to open up the LAP to liberate active TGF-β1^19^, passive remodeling force may not be or at least do not happen at an appreciable occurrence to significantly change myofibroblast activation in established microtissues. On the other hand, external stretching that simulates breathing in the lung and pulsatile flow in vasculature is capable of maintaining myofibroblast behavior in microtissues through this autocrine-signaling pathway.

## Conclusions

In this article we demonstrated the effectiveness of MVAS-force devices to investigate fibrosis by showing that microtissues are able to recapitulate characteristic mechanical and biological features that occur in fibrotic disease, namely increased contractility, stiffness and α-SMA expression. These changes were partially reversible, upon exogenous TGF-β1 withdrawal. In addition to its ability for relatively high throughput mechanical assessment of tissue-level mechanics, the MVAS-force permitted long-term conditioning of fibrotic microtissues to mimic the stretch cells experience in lung and vasculature tissue. Importantly stretch inhibited the partial myofibroblast deactivation that occurred following exogenous TGF-β1 withdrawal likely through endogenous latent TGF-β1 activation. Therefore external mechanical stretch might be a powerful stimulus for continued myofibroblast activation to progress fibrotic development. To conclude, further research into this pathway, which links mechanical forces to soluble mediators, may have important implications in the development of therapeutic treatments urgently needed to stop the advancement of fibrosis.

## Additional Information

### Conflicts of Interest

There are no conflicts to declare.

## Acknowledgements

M.W. is supported by OGS (Ontario Graduate Scholarship). The authors acknowledge support from individual NSERC Discovery Grants (M.G. and A.E.P.). M.G. and A.E.P. acknowledge the Canadian Foundation for Innovation.

## Data Availability

The data generated during the current study is available from the corresponding author upon reasonable request.

