## Supplementary information for "Mechanical stretch sustains myofibroblast phenotype and function in microtissues through latent TGF-β1 activation"

*Condensed Running Title: TGF-β1 activation in microtissues*

Matthew Walker[1], Michel Godin[2,3,4], Andrew E. Pelling[1,2,5,6]*

[1]Department of Biology, Gendron Hall, 30 Marie Curie, University of Ottawa, Ottawa, ON, K1N5N5 Canada
[2]Department of Physics, 150 Louis Pasteur pvt., STEM Complex, University of Ottawa, Ottawa, ON K1N 6N5 Canada
[3]Department of Mechanical Engineering, Colonel By Hall, 161 Louis Pasteur, University of Ottawa, Ottawa, ON K1N6N5 Canada
[4]Ottawa-Carleton Institute for Biomedical Engineering, Colonel By Hall, 161 Louis Pasteur, University of Ottawa, Ottawa, ON K1N6N5 Canada
[5]Institute for Science Society and Policy, Simard Hall, 60 University, University of Ottawa, Ottawa, ON, K1N5N5 Canada
[6]SymbioticA, School of Human Sciences, University of Western Australia, Perth, WA, 6009 Australia

*Keywords:*
Myofibroblasts, Fibrosis, Microtissue, TGF-β1, Stretch, Microfabrication

* Author for correspondence
Andrew E. Pelling
150 Louis Pasteur pvt.
University of Ottawa
Ottawa, ON K1N 6N5
Canada

Web: http://www.pellinglab.net

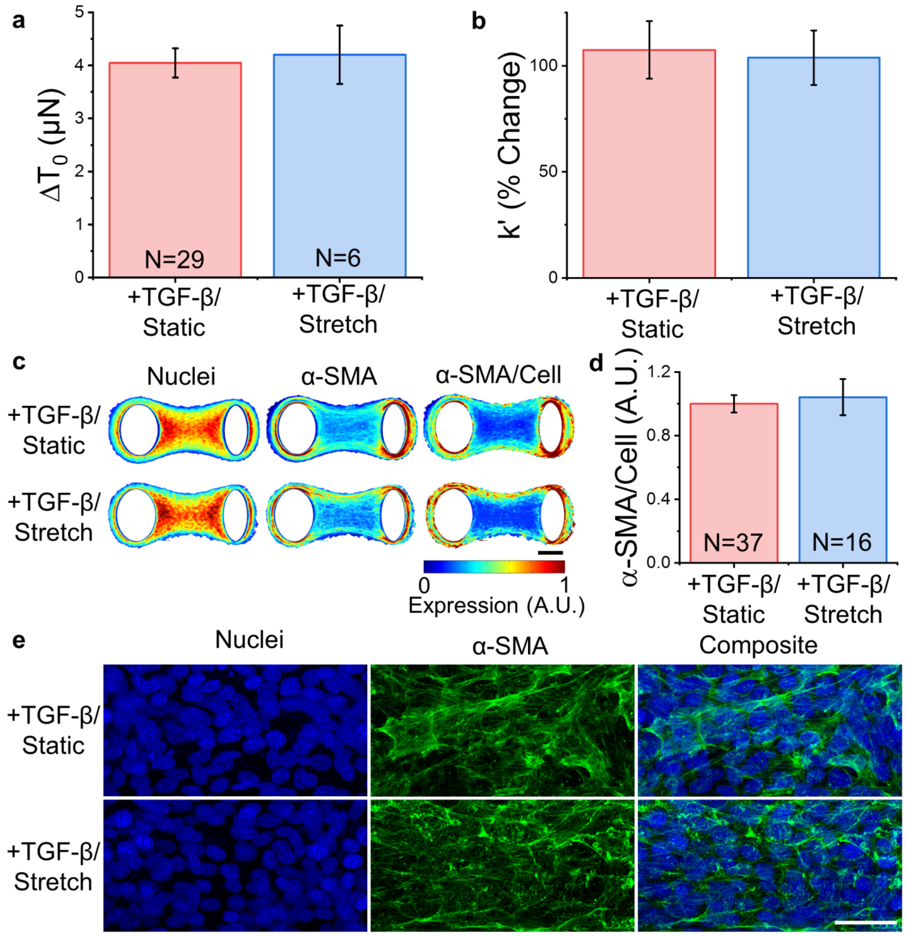


**SI 1: Cyclic stretch does not contribute to myofibroblast differentiation in microtissues.** Stretching microtissues during TGF-β treatment did not change contractility (a), stiffness (b) or α-SMA expression (c-e) compared to a static control group (t-tests, P>0.05).
